# Incidences of Helicobacter infection in pigs and tracing occupational hazard in pig farmers

**DOI:** 10.1101/2023.03.27.534128

**Authors:** Seema Rani Pegu, Swaraj Rajkhowa, Pranab Jyoti Das, Joyshikh Sonowal, Gyanendra Singh Sengar, Rajib Deb, Manjisa Choudhury, Nabajyoti Deka, Souvik Paul, Juwar Doley, Dilip Kumar Sarma, Samir Das, N. H. Mohan, Rajendran Thomas, V.K. Gupta

**Affiliations:** ICAR- National Research Centre on Pig, Rani, Guwahati, 781131, Assam; College of Veterinary Science, AAU, Khanapara, Guwahati, 781022, Assam; ICAR Research Complex for NEH Region, Umiam, 793103, Meghalaya

**Author notes:** Corresponding Author:**Seema Rani Pegu**-Senior Scientist (Veterinary Pathology), ICAR-National Research Centre on Pig, Rani, Guwahati, Assam, India, **Pranab Jyoti Das**-Principal Scientist (Animal Genetics and Breeding), ICAR-National Research Centre on Pig, Rani, Guwahati, Assam, India.

**Keywords:** *Helicobacter Suis*, *Helicobacter Pylori*, Urease Test, Gastric Ulcer, Pathology, non-*Helicobacter pylori* Helicobacter (NHPH)

## Abstract

*Helicobacter species* (*H. sp*.) is a gram-negative spiral-shaped motile bacteria that causes gastritis in pigs and also colonizes the human stomach. The current study seeks to assess the prevalence of various *H. sp*. in the gastric mucosa of slaughtered and dead pigs, as well as the prevalence of *Helicobacter* infection among pig farmers. A total of 403 stomach samples from various pig slaughter points, 74 necropsy samples from various pig farms and 97 stool samples from pig farmers were collected from Assam, India. Among 477 pig stomach samples tested, 214 samples with gastritis (20.09%) showed Gram negative, spiral-shaped organisms in brush cytology from the mucosal surface, and the rest of the 263 stomach samples without any gastric lesion showed only 3.04% Gram negative, spiral-shaped organisms. In ultrastructure investigation, Scanning Electron Microscopy (SEM) of the four urease positive stomach samples revealed a tightly coiled *Helicobacter bacterium* (spiral-shaped) found in the mucous lining of the stomach. In histopathological examination of pars esophagia, cardiac and fundic mucosa showed chronic gastritis associated with hemorrhagic necrosis, leucocytic infiltration with neutrophils and macrophages, and lymphoid aggregates (lymphoid follicles) etc. PCR confirmed 16S rRNA genes of *Helicobacter suis* (*H. suis*) where a total of 42 (19.63%) out of 214 pig stomach samples and 2 (2.08%) out of 96 stool samples of pig farmers were found positive for *H. suis*. of these 96 stool samples of pig farmers 3 (3.12%) were confirmed positive for *Helicobacter pylori* (*H. pylori*) Phosphoglucosamine mutase gene in PCR. Phylogenic analysis of the 16S rRNA gene of *H. suis* showed distinct clusters with other *H*. sp. In conclusion, this study provides evidence for the prevalence of *Helicobacter* both in pig gastric mucosa and human stool. The findings highlight the need for improved sanitation and hygiene practices among pig farmers to minimize the risk of *Helicobacter* infection in humans.

## 1. Introduction

Ulceration of gastric mucosa is a commonly occurring ailment in pigs worldwide (Haesebrouck *et al*., 2009). Although seldom manifested clinically, the condition is responsible for substantial economic losses on account of decreased feed intake and decreased average daily weight gain (De Witte *et al*., 2017). Mucosal changes due to ulceration may vary from mild epithelial changes to ulcers extending to the entire glandless region of the stomach at later stages. The etiology of gastric ulceration in pigs is multifactorial, including pelleting of feed, fine particle sized feed, presence of highly fermentable carbohydrates as well as short-chain fatty acids in feed, interruption in feeding, overcrowding/ stress, respiratory ailments, *H. suis* infection, etc (De Bruyne *et al*., 2012). The pathogenesis of gastric ulcers in pigs remains still unclear, the ulcers are mostly confined to the pars oesophagea, a small glandless area situated just near the opening of the oesophagus (Haesebrouck et al., 2009). This area is glandless and therefore not protected by mucous against the corrosive action of hydrochloric acid produced in the fundic region of the stomach, and this repeated exposure of the area to acid may result in ulceration (De Witte et al., 2017). *H. suis*, a common zoonotic enteropathogen of pigs inhabit the fundic and pyloric region of gastric mucosa. The alteration of gastric environment due to colonization of *H. suis* has been suggested to manipulate the hydrochloric acid production and thereby play an important role in the development of gastric ulceration in pigs (Hellemans *et al*., 2007, Herskin *et al*., 2016). *H. suis* has a particular affinity for acid producing parietal cells and in histological sections has been found near or even inside the canaliculi of parietal cells (Hellemans et al., 2007). *H. suis* has been incriminated to disturbing the delicate gastric equilibrium in either of many ways, by inciting necrosis and associated degenerative changes in parietal cells (Joo *et al*., 2007, Flahou *et al*., 2010), by affecting the expression of genes regulating the gastric hydrogen potassium ATPase proton pump in parietal cells (Zhang *et al*., 2016) or by modification of the function of G-cells producing gastrin or D-cells producing somatostatin (Sapierzyński *et al*., 2007).

*H. sp*. are considered to be one of the predominant pathogens associated with a wide range of gastric ailments in humans(Joosten *et al*., 2016). Besides *H. pylori, H. suis* is the most prevalent gastric non-*H. pylori Helicobacter* (NHPH) species found in patients with gastric disease (Van den Bulck *et al*., 2005). *H. suis* is an important zoonotic organism (Joosten *et al*., 2013) and several reports suggest that pigs are the source of *H. suis* infection in human beings (Flahou *et al*., 2018). NHPH infection has been found to be associated with gastric diseases, such as peptic ulcers, chronic gastritis, gastric cancers, and gastric mucosa-associated lymphoid tissue lymphoma (Debongnie *et al*., 1998, Morgner *et al*., 1995, Morgner *et al*., 2000). *H. suis* is frequently detected in adult patients, whereas infection of *H. pylori* is very rare in adults (Mendall *et al*., 1992), moreover sometimes *H. suis* is detected in patients after H. pylori eradication therapy (Blaecher *et al*., 2013) emphasizing the role of NHPH as an important human gastric pathogen.

Despite *H. suis* being an important pathogen in pigs and humans, no reports have been published on prevalence of *H. suis* among pigs and zoonotic potential of *H. suis* among pig farmers in India. Therefore, the present study was undertaken to evaluate the prevalence of different *H. sp*. in the gastric mucosa of slaughtered as well as dead pigs (particularly in cases of gastritis) by various diagnostic assays and also to evaluate the incidence of *Helicobacter* infection among the pig farmers.

## 2. Materials and Methods

### 2.1. Tissue and clinical samples

A total of 477 gastric tissue samples from slaughtered pigs (n=403) were collected from different slaughter points and (n=74) necropsy samples from different pig farms of Assam, India were collected during the study. In addition to this and 97 nos. of stool samples were collected from pig farmers and pig handlers to study the incidence of *Helicobacter* infection. All the samples were analyzed through urease test and PCR assay.

### 2.2. Urease Test

#### 2.2.1. Rapid Urease Test (RUT) dry test

The gastric mucosal samples were screened by a commercial rapid urease kit i.e. RUT DRY Test kit as per the test protocol instruction (Gastro Cure System, Kolkata, India). In nutshell, test samples were directly put into the well containing urea and the urease enzyme produced by *H. sp*. rapidly hydrolyses the urea in the well, producing ammonia. The rise in the pH of the medium by ammonium ions was detected with a pH indicator. The test was read after 3 hours for positive and negative results.

#### 2.2.2. Modified Rapid Urease test

A modified rapid urease test (Kolodzieyski *et al*., 2008) was performed on the small mucosal specimens cut out with scissors from four different parts of stomach (namely -par esophagia, cardiac, fundic and pyloric region) and placed in test tubes with two drops of physiological solution (distilled water) during the collection time. The tubes were brought to the laboratory; the solution was removed followed by transferring of each mucosal specimen to each well of a 96-well plate. 1 ml of reagent containing 10% unbuffered urea, pH 6.8 and 1% phenol red was added to every sample. The results were recorded at 30 and 60 min after addition of the reagent. A change of colour from pale yellow to bright pink was considered a positive reaction.

### 2.3. Cytological examination

Mucosal scrapings were collected from the gastric mucosa using cotton buds. The buds were rolled over the mucosa at the sample site and subsequently rolled onto a clean slide. The preparations were then air dried, Gram’s stained and examined using an oil immersion lens at 100 X magnification.

### 2.4. Gross and Histopathological examination

Stomach tissues with gastritis and ulcerative lesions were subjected for gross as well as microscopic examination to record the pathological changes in the stomach. The stomach tissue samples from different regions (par esophagia, cardiac, fundic, and pyloric) were collected and preserved in 10% formalin solution for routine histopathological examination. After proper fixation the tissues were processed, embedded in paraffin and 4–5 μ thick sections were made, stained with Haematoxylin and Eosin for histopathological studies.

### 2.5. Isolation of DNA from stomach samples and stool samples

DNA was extracted from gastric mucosa of pigs and stool sample of human using the DNeasy Tissue Kit (QIAZEN, Germany) and QIAampStoolDNA Mini Kit (QIAZEN, Germany) respectively, according to the manufacturer’s instruction. The extracted DNA was stored at -20^0^C until further analysis. Qualitative and quantitative assessment was done by agarose gel electrophoresis (0.8%) and nanodrop readings using eppendorfBioPhotometer plus at optical density (OD) 260nm/280nm, respectively.

### 2.6. Polymerase Chain Reaction (PCR) assay

A set of primers were used for detecting two *H. sp*. Such as *H. suis* and *H. Pylori* (Supplementary Table 1) as per Proietti *et al*. (Proietti *et al*., 2010). The amplified products were run on 1.5% agarose gel and photographed under UV illuminator. Negative controls were used for the validation of the PCR.

### 2.7. Phylogenetic analysis

Phylogenetic analysis of Sanger sequenced four 16S rRNA gene of *H. sius* with 12 reference sequences and two phosphoglucoseamine mutase gene of *H. pylori* with 8 reference sequences were done separately. The reference sequences were retrieved from the NCBI GenBank nucleotide database (Supplementary Table 2). Phylogenetic and molecular evolutionary analyses were conducted using DNASTAR software with the Neighbor-Joining Method.

### 2.8. Scanning Electron Microscopy (SEM)

Stomach samples collected from dead piglets during post mortem and which were subsequently tested positive by RUT dry test, were selected for SEM analysis. The tissues were fixed in 2.5% Glutaraldehyde and 0.1M Cacodylate buffer and subjected for SEM at SAIF, NEHU Shillong, Meghalaya.

## 3. Results

### 3.1. Urease activity in RUT dry test

Urease activity was monitored by observing the immediate changes in colour, pink colour in case of positive samples while in case of negative, the well remained yellow (Fig. 1(a), 1(b)). Urease activity was observed in 119 (24.94%) of the 477 gastric samples from pigs. Out of 119 samples found to be positive for urease activity, 112 samples were associated with gross gastric lesions and the others were not. A total of 11 (11.45%) out of 96 fecal samples of pig farmers were found to be urease positive.

**Fig. 1:**
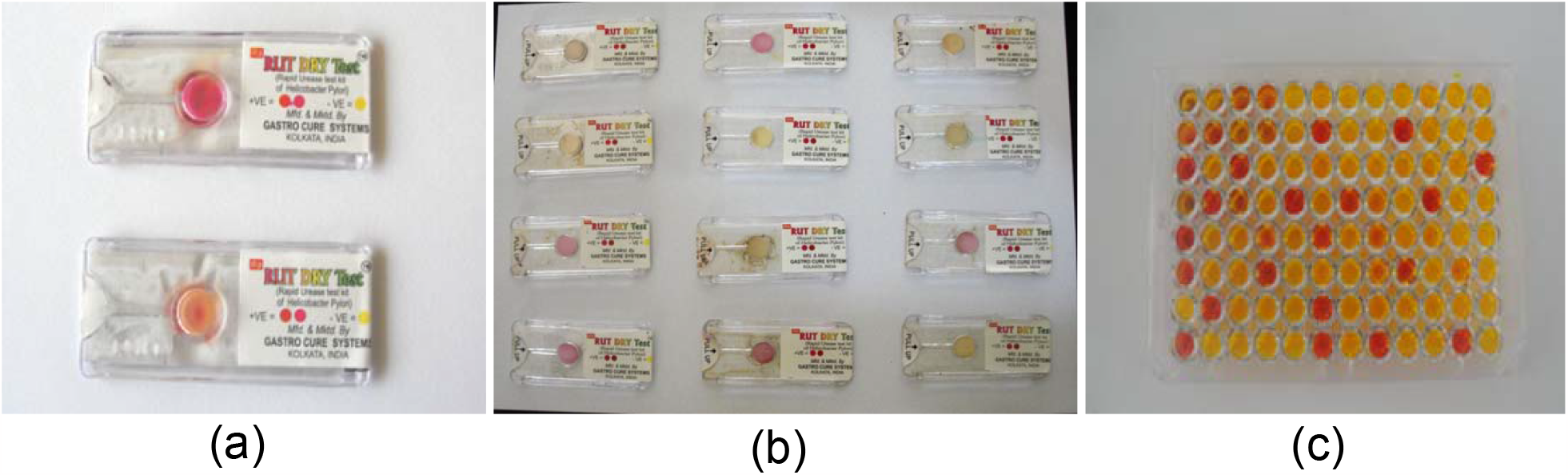
Urease test of stomach sample of pig. Here, (a) Urease enzyme positive gastric mucosal samples of pigs in RUT dry test. (b) Urease enzyme positive fecal samples of pig farmers in RUT dry test. (c) Modified Rapid Urease Test in 96-well plate detected the urease enzyme of *Helicobacter* in gastric mucosal samples of slaughtered and necropsied pigs with gastritis. Pink showing positive and yellow showing negative results.

### 3.2. Urease activity in Modified Rapid Urease Test in 96-well plate

A change of colour from pale yellow to bright pink was observed in positive samples in contrast to the pale yellow colour observed in negative samples (Fig. 1(c)). Among the 112 numbers of urease positive sections of the stomach, the urease activity was observed in the pars esophagia number (33 %) and cardiac number (40.17%) were tested highly positive in comparison to fundic number (16.07%) and pyloric number (8.92%) sections in the 96-well plate.

### 3.3. Cytological examination

Among 477 numbers of pig stomach samples, 214 samples with gastritis (20.09%) showed Gram negative, spiral shaped organisms in brush cytology from the mucosal surface (Fig. 2). Rest 263 stomach samples without any gastric lesion showed only 3.04% Gram negative, spiral shaped organisms. These organisms were detected mainly in the cardiac, fundic and par oesophageal part of the stomach.

**Fig. 2:**
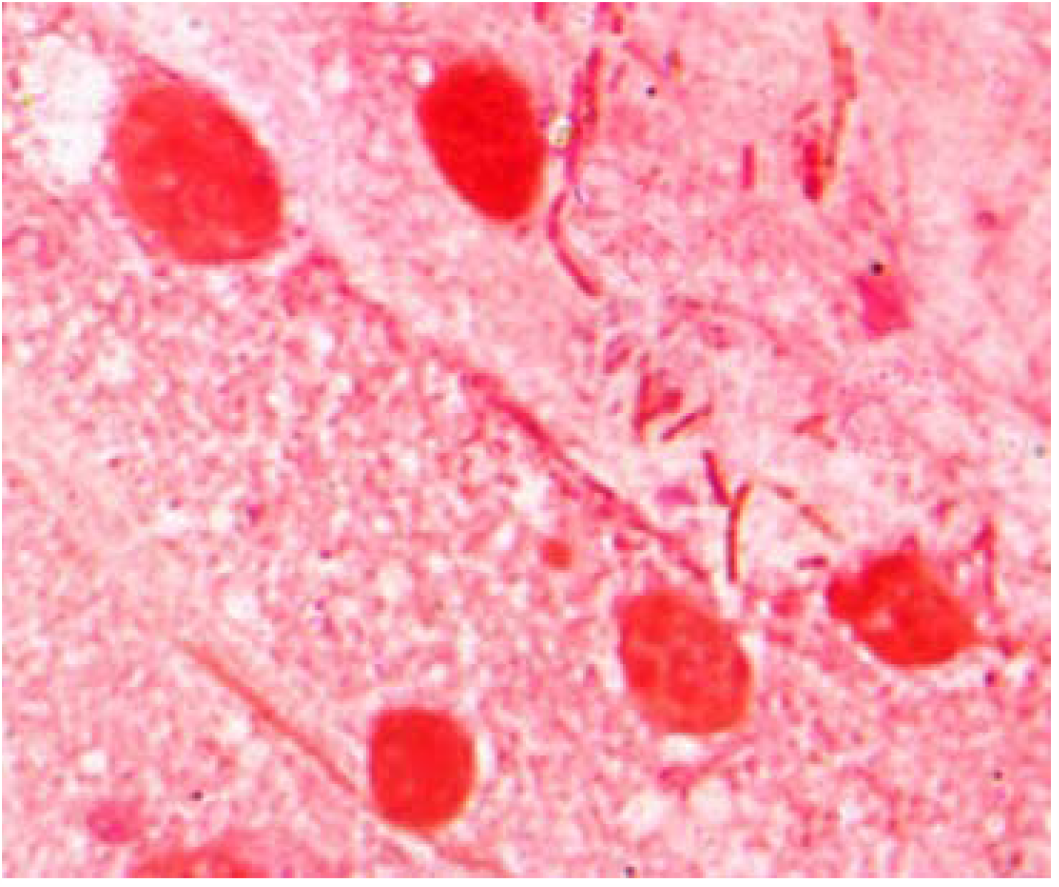
Cytological examination followed by Gram staining showing tightly spiraled bacteria in the gastric mucosa of a pig (Gram’s staining X 100 x)

### 3.4. Gross Pathological study

Ulcerative lesion in the pars esophagia and cardiac region of the stomach (Fig. 3a) were recorded in the slaughtered animals while mild to severe gastritis were recorded in the necropsied pigs. (Fig. 3b). In the present study we have observed 214 cases of mild to severe gastritis whereas 49 cases of gastric ulcers during gross pathological examination of the total 477 pig stomach samples.

**Fig. 3:**
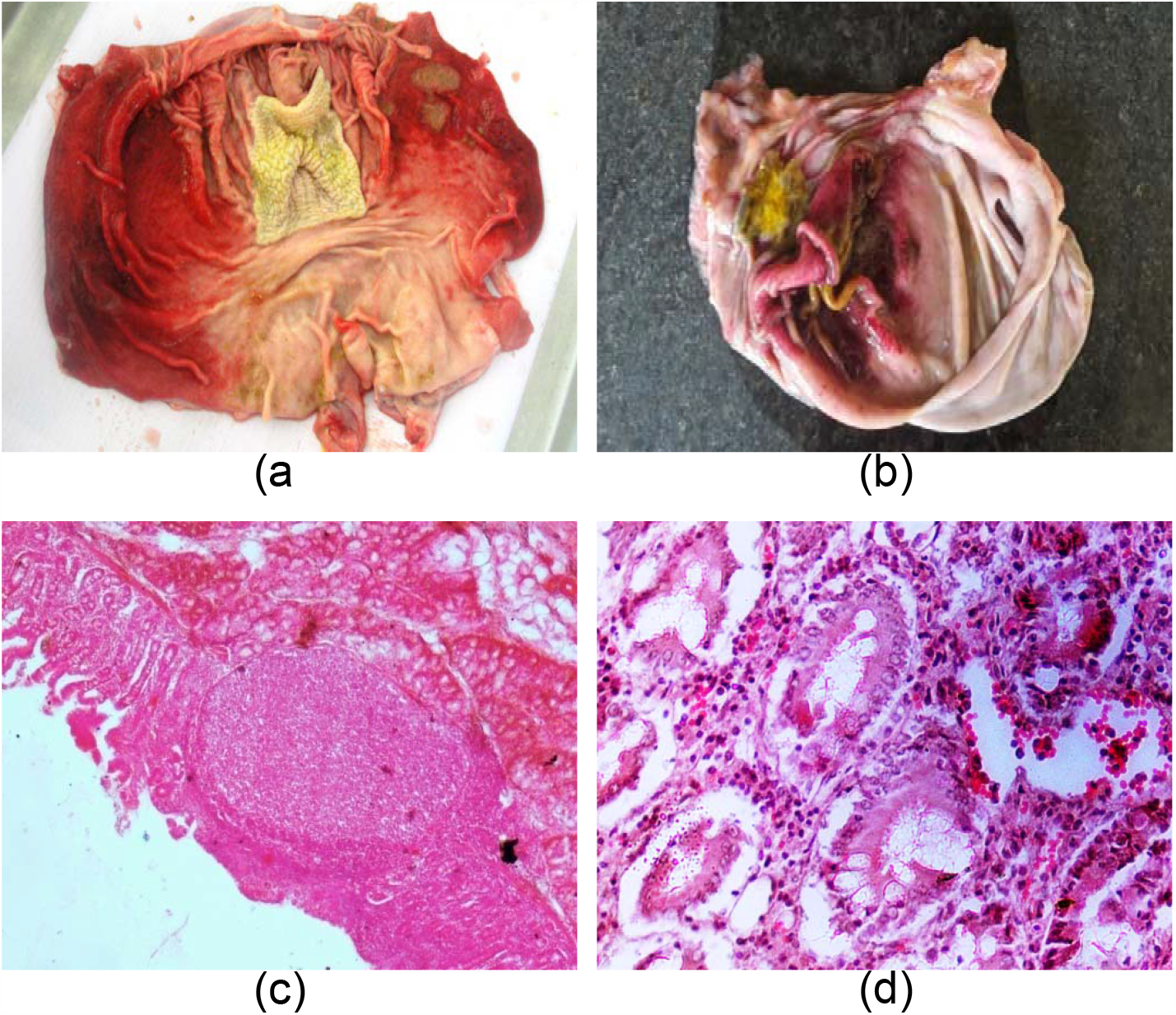
Gross and histopathological lesions of pig stomach. Here, (a) Ulcerative lesion in the cardiac part of Gastric mucosa in a slaughtered pig, (b) Gastric lesion in the cardiac region of gastric mucosa in a necropsied pig, (c) Lymphoid follicle in the cardiac gastric mucosa of a *Helicobacter* infected pig (H & E X 4x). (d) Fundic mucosa of pigs with gastritis (H & E X 20x).

### 3.5. Histopathological studies

Histopathological examination revealed pars esophagia, cardiac and fundic mucosa of pig with chronic gastritis (Fig. 3d). Other histopathological findings included necrosis with haemorrhage, leucocytic infiltration with neutrophils and macrophages as the main cell types. Lymphoid aggregates (Lymphoid follicle) was one of the major lesions observed in the gastric mucosa of stomach samples and owing to the size of the follicles, they were associated with displacement and loss of gastric glands (Fig. 3c).

### 3.6. Molecular detection of *Helicobacter* by PCR

PCR confirmed 16S rRNA genes of *H. suis* where a total of 42 (19.63%) out of 214 pig stomach samples and 2 (2.08%) out of 96 stool samples of pig farmers were found positive for *H. suis* (Fig. 4a). of these 96 stool samples of pig farmers 3 (3.12%) were confirmed positive for *H. pylori* Phosphoglucosamine mutase gene in PCR. (Fig. 4b). However, none of the pig stomach sample was detected positive for *H. pylori* in PCR analysis.

**Fig. 4:**
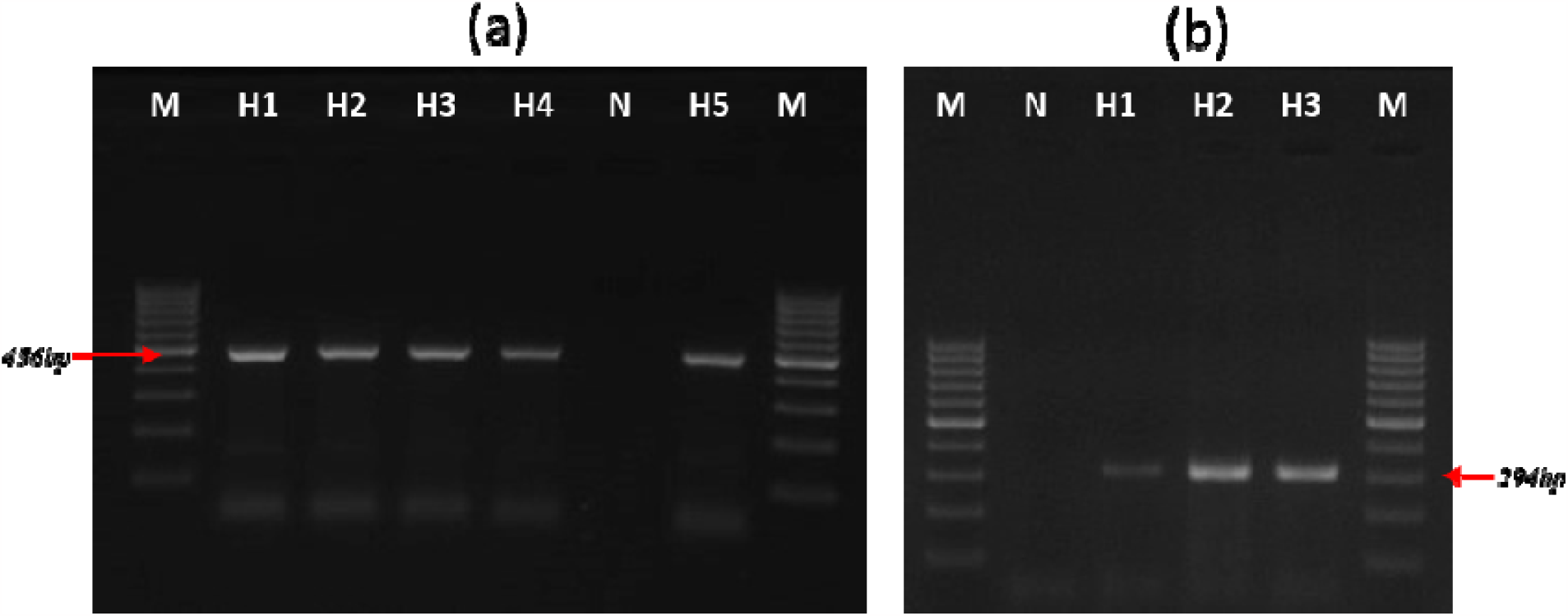
PCR detection of *Helicobacter species* (a) here, M:100bp DNA marker, H1, H2, H3: pig stomach sample, H4, H5: human fecal sample, N: Negative control. H1 (Positive control), H2, H3, H4, H5 showed positive for *Helicobacter suis* (456bp). (b) here, M :100bp DNA marker, H1, H2, H3: Human fecal sample, N: Negative control. H1, H2, H3 showed positive for *Helicobacter pylori* (294bp).

### 3.7. Phylogenetic analysis

Sequence analysis of 16s RNA gene of pig gastric mucosa samples and stool samples of human with 18 other *H. sp*. sequences retrieved from the NCBI database showed distinct cluster of *H. suis*. The *Helicobacter* strains in the present study i.e. *Helicobacter suis* _Assam_ Human1 and *Helicobacter suis*_Assam_Pig1 showed 99.8% similarity with each other, and pairwise distance identity of this two sequences showed the highest identity of 99.6 to 99.8% with the study sequences *Helicobacter suis*_Assam_Human2, *Helicobacter suis*_Assam_Pig2 and also with the reference sequence Candidatus *Helicobacter suis*1 and Candidatus *Helicobacter suis*2 (Fig. 5). The phosphoglucoseamine mutase gene sequence analysis of two *H. pylori* positive fecal samples of pig farmers in the present study i.e. OQ409904 and OQ409903 showed highest similarity of 98.3% with the reference sequence of *H pylori* retrieved from NCBI (CP024948 and CP0202071) and (Fig. 6).

**Fig. 5:**
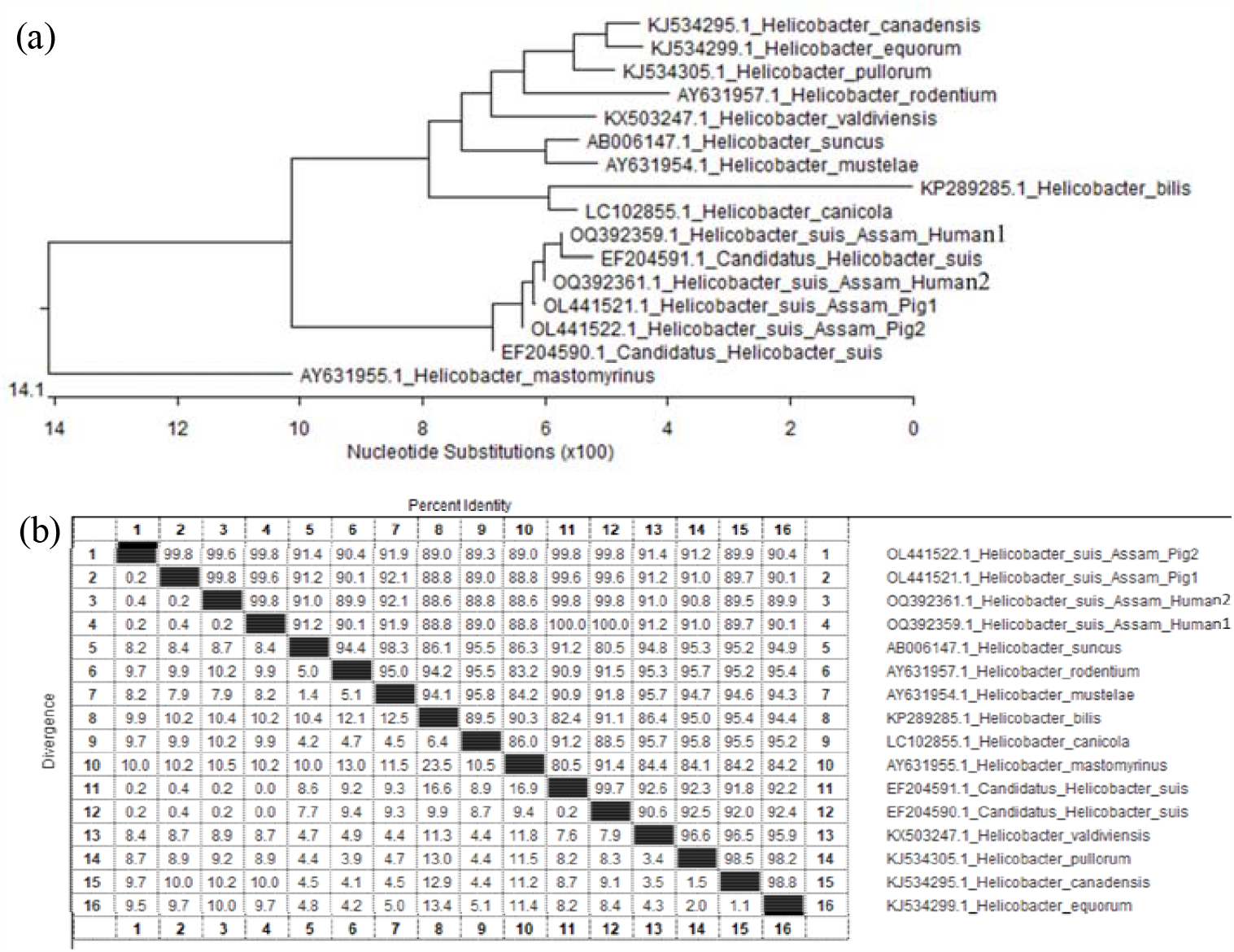
Phylogenetic tree of *Helicobacter suis* from pig stomach samples. (a) Sequence analysis of *Helicobacter suis* PCR positive fecal samples of pig with other sequences of *Helicobacter sp*. And Pairwise distance analysis of 16s rRNA gene of *Helicobacter sp*. from pig and human samples.

**Fig. 6:**
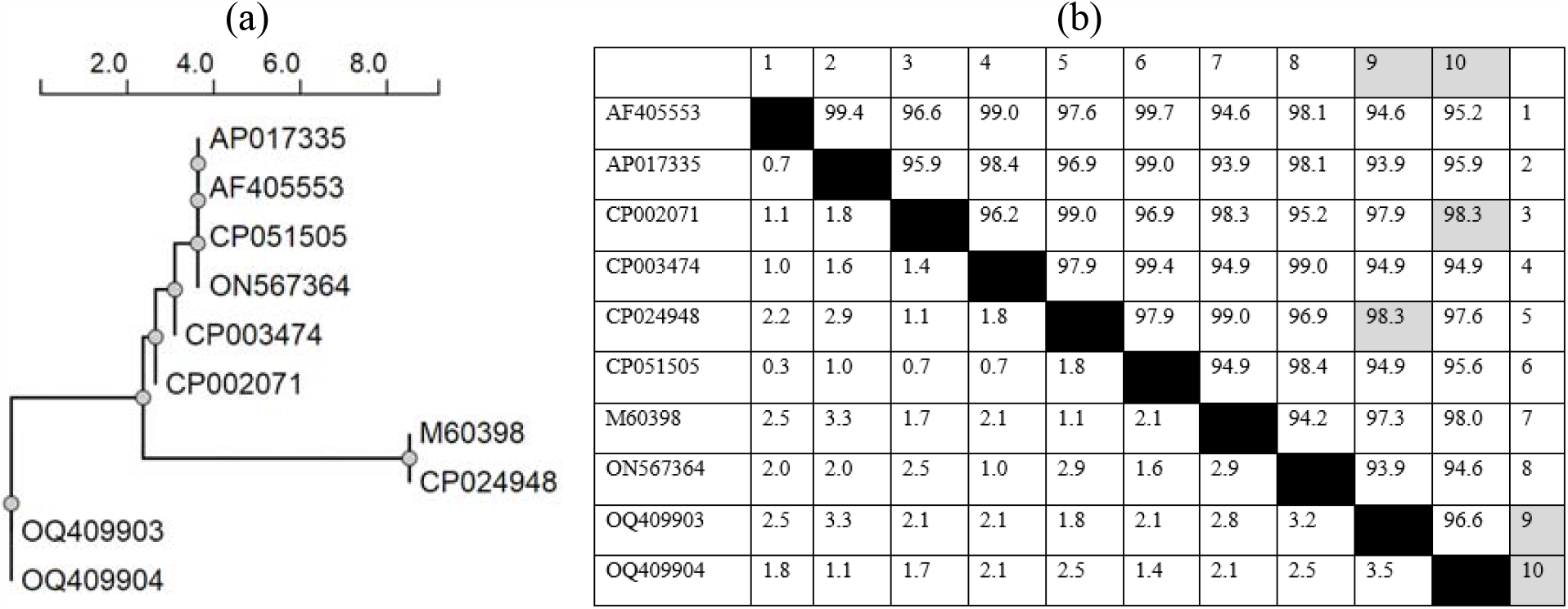
Phylogenetic tree of *Helicobacter pylori* from human fecal samples. (a) Sequence analysis of *Helicobacter pylori* PCR positive fecal samples of human revealed identity with phosphoglucosamine mutase gene of *Helicobacter pylori* (b) Pairwise distance analysis of *Helicobacter pylori* from human samples showing a maximum identity of 98.3%.

### 3.8. Scanning electron microscopy

Scanning Electron Microscopy of the four urease positive stomach samples revealed tightly coiled *Helicobacter* bacterium (spiral-shaped) found in the mucous lining of the stomach (Fig. 7).

**Fig. 7:**
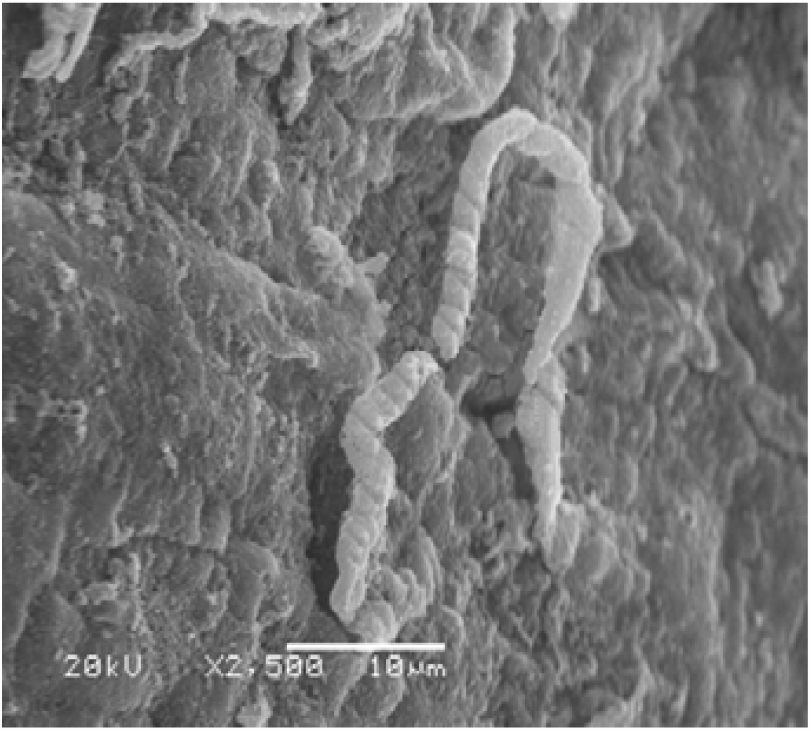
*Helicobacter* bacterium (spiral-shaped) found in the mucous lining of the stomach (SEM)

## 4. Discussion

*Helicobacter* like organisms in stomach of pigs were first encountered during histological studies (Queiroz *et al*., 1990). Thereafter various studies carried out through the world have revealed that *H. suis* infection among pigs are common, with prevalence rate ranging above 50% (De Witte et al., 2017, Flahou et al., 2018). Diverse range of nonspecific diagnostic methods along with latest diagnostic tools have been used for the detection of the organism in biological samples (Kolodzieyski et al., 2008). Under the current study we used rapid urease test (RUT) dry test, modified rapid urease test, cytological and histopathological studies, PCR and SEM for detection of *Helicobacter* infection among pig and occupational pig handlers/rearers.

In the present study the incidence of *Helicobacter* infection among pigs ranged between 20-25%, as per the results of the different detection methods. There are no reports of *Helicobacter* infection among pigs are available from India, however the incidence rate was in line with the reports from other parts of world (De Witte et al., 2017). The notably high prevalence in adult pigs suggested that the immune system was not able to get rid of the infection. It has also been suggested that *H. suis* imposes some sort of immune suppressive effect on the host by down regulating expression of few cytokines (Bosschem *et al*., 2017).

Erosion and gastric ulceration in the lining of stomach especially in the non-glandular oesophageal portion among the slaughtered pigs is a common condition (Friendship, 1999, Friendship & production, 2004). The causes of gastric ulceration are multifactorial including genetic, nutritional, environmental and infectious agents (De Witte *et al*., 2018). Associations have been made between *Helicobacter* infection and the presence of ulcers in the pars oesophageal of the stomach (Haesebrouck et al., 2009). In the present study we have observed 214 cases of mild to severe gastritis and 49 cases of gastric ulcers by gross and histopathological analysis.

Since all gastric *H. sp*. produce urease enzyme, hence the rapid urease - producing property has been utilized to detect the bacterium with a urease test. This urease test has an advantage of the test result interpretation within an hour. The percentage of detection of positive results determined by the urease test (24.94%) and cytological examination (20.1%). In the present study it was observed that the percentage positivity of urease and cytological examination was directly proportional to the gastric lesion recorded in the stomach.

Gross pathological alteration recorded were mostly varying degrees of gastritis with ulcerative lesion mainly in the par oesophageal and cardiac region. Similar findings were also recorded by the earlier workers (Szeredi *et al*., 2005). Histopathological studies revealed lymphoid aggregates associated with displacement and loss of gastric glands, as reported elsewhere (Hellemans et al., 2007, Joosten et al., 2013).

PCR confirmed the 16S rRNA genes of *H. suis*, where a total of 42 (19.63%) out of 214 pig stomach samples and 2 (2.08%) out of 96 stool samples from pig farmers were found positive for *H. suis*. Of these 96 stool samples from pig farmers, three (3.1%) were confirmed positive for the *H. pylori* phosphoglucosamine mutase gene by PCR. Recently, *H. pylori* and *H. suis* were reported in free-roaming wild boars (*Sus scrofa*) (Teixeira *et al*., 2021), indicating that these microbes have close contact with domestic animals, wildlife, and people, increasing the threat of bacterial transmission across species. *H. suis* is also prevalent in non-human primate *Macacamulatta* where in some cases clinical importance remains unclear (Haesebrouck et al., 2009, Marini *et al*., 2021).

Among the samples collected from the occupational pig farmer, 11 out of 96 were urease positive, and PCR positivity was around 2.08%. Two nos. of samples were found to be positive for *H. suis* and three samples were *H. pylori* in stool samples collected from pig farmers. So in a subclinical condition, this is an indication of the prevalence of *H. suis* and *H. pylori among pig farmers*. Our study suggesting that *Helicobacter* infection was a notable occupational hazard among the pig farmers. Researchers (Yasuda *et al*., 2022) from Japan reported PCR detection for non-*H. pylori Helicobacter* (NHPH) in 30 cases, 26 of which were found positive as *H. suis* and 2 as *Helicobacter heilmanii*/*Helicobacter ailurogastricus*. Further study is required to draw a possible association between the zoonotic transmissions of the disease from pig to pig farmers. In terms of occupational hazards, studies have suggested that pig farmers and individuals who work closely with pigs may be at an increased risk of Helicobacter infection. The risk of infection may vary depending on the specific farming practices and biosecurity measures in place. Overall, while Helicobacter infection in pigs is not typically a significant health risk for the animals themselves, it is important for individuals who work closely with pigs to take appropriate precautions to minimize the risk of infection. This may include wearing appropriate protective clothing and practicing good hygiene, such as washing hands regularly and avoiding touching the face or mouth while working with pigs.

## 4. Conclusion

This study provides the first insights into the evidence and phylogeny of NHPH from the different regions of Assam, India. The evidence shows a possibility of transmission of *H. spp*. from the pig population to humans. Although the Helicobacterium is a rare pathogenic organism for swine, it is a major concern in respect of occupational health workers associated with their livelihood in piggery husbandry practices.

## Acknowledgement

The authors are thankful to the Director, ICAR-National Research Center on Pig, Guwahati, Assam, India for providing the necessary facilities to carry out the present research.

## Authors contribution

Concept and Designing of the manuscript: SRP. Wet-lab work: SRP, MC, NJD, SP, RD, and JD. Dry lab work: PJD and JS. Manuscript drafting: SRP, PJD, JS and RD. Proofreading and editing: GSS, SR, DKS, and VKG. Resources and supervision: SRP, SR, NHM, RT. All authors revised and approved the final version of the manuscript.

## Data Availability Statement

The data that support this study will be shared upon reasonable request to the corresponding author.

