## Supplementary table 1 and Supplementary table 2 for "Incidences of Helicobacter infection in pigs and tracing occupational hazard in pig farmers"

**Supplementary Tables:**

| **Supplementary Table 1: The set of primers used for PCR assay** | | | | |
| --- | --- | --- | --- | --- |
| **Species** | **Gene** | **Oligonucleotides and sequence (5’ para 3’)** | **Product Size** | **Reference** |
| *H. suis* | 16S rRNA | F : TTGGGAGGCTTTGTCTTTCCA | 433bp Proietti *et al*. (2010)  or  456bp (from current study) | |
|  |  | R: GATTAGCTCTGCCTCGCGGCT |  |  |
| *H. pylori* | Phosphoglucosamine mutase | F : AAGCTTTTAGGGGTGTTAGGGGTTT | 294bp | Proietti *et al*. (2010) |
|  |  | R : AAGCTTACTTTCTAACACTAACGC |  |  |

| **Supplementary Table 2**: **Gene bank accession numbers of reference sequences used for the present study** | | |
| --- | --- | --- |
| **Sl. No.** | **GenBank Accession No.** | **Reference Sequence** |
| 1. | KJ534299.1 | *Helicobacter equorum* |
| 2. | KJ534295.1 | *Helicobacter canadensis* |
| 3. | KJ534305.1 | *Helicobacter pulloram* |
| 4. | KX503247.1 | *Helicobacter valdiviensis* |
| 5. | EF204590.1 | *Candidatus helicobacter suis* |
| 6. | EF204591.1 | *Candidatus helicobacter suis* |
| 7. | AY631955.1 | *Helicobacter mastomyrinus* |
| 8. | LC102855.1 | *Helicobacter canicola* |
| 9. | KP289285.1 | *Helicobacter bilis* |
| 10. | AY631954.1 | *Helicobacter mustelae* |
| 11. | AY631957.1 | *Helicobacter rodentium* |
| 12. | AB006147.1 | *Helicobacter suncus* |
| 13. | OQ392359.1 (This study) | *Helicobacter suis*_Assam_Human1 |
| 14. | OQ392361.1 (This study) | *Helicobacter suis*_Assam_Human2 |
| 15. | OL441521.1 (This study) | *Helicobacter suis*_Assam_Pig1 |
| 16. | OL441522.1 (This study) | *Helicobacter suis*_Assam_Pig2 |
| 17. | OQ409903 (This study) | *Helicobacter pylori_India* |
| 18. | OQ409904 (This study) | *Helicobacter pylori_India* |
| 19. | AP017335.1 | *Helicobacter pylori_Japan* |
| 20. | AF405553.1 | *Helicobacter pylori_USA* |
| 21. | CP051505.1 | *Helicobacter pylori_Peru* |
| 22. | ON567364.1 | *Helicobacter pylori_Iraq* |
| 23. | CP003474.1 | *Helicobacter pylori_Peru* |
| 24. | CP002071.1 | *Helicobacter pylori_Peru* |
| 25. | M60398.1 | *Helicobacter pylori_France* |
| 26. | CP024948.1 | *Helicobacter pylori_South Korea* |
